# Modelling donor screening strategies to reduce the risk of SARS-CoV-2 via fecal microbiota transplantation

**DOI:** 10.1101/2020.06.24.169094

**Authors:** Scott W. Olesen, Amanda Zaman, Majdi Osman, Bharat Ramakrishna

**Affiliations:** OpenBiome, 2067 Massachusetts Avenue, Cambridge, MA 02140

## Abstract

The potential for transmission of SARS-CoV-2 shed in stool via fecal microbiota transplantation is not yet known, and the effectiveness of various testing strategies to prevent FMT-based transmission has also not yet been quantified. Here we use a mathematical model to simulate the utility of different testing strategies.

## INTRODUCTION

Fecal microbiota transplantation (FMT), the instillation of stool from a healthy donor into a patient’s gut, is a recommended therapy for the most common hospital-acquired infection in the United States, *Clostridioides difficile*, and is being explored as an experimental therapy for dozens of other conditions.^1,2^ As with all human-derived therapies, the safety of FMT depends on screening donors to prevent transmission of pathogens via the procedure,^3^ and screening guidelines must be continually updated to account for emerging pathogens.

SARS-CoV-2, the virus that causes COVID-19, is primarily considered a respiratory pathogen, but evidence suggests that the virus is able to independently replicate in the gut, raising the possibility of transmission via the fecal oral-route or via FMT.^4^ Practitioners^4,11^ and regulators^12^ have therefore called for screening of FMT donors for SARS-CoV-2. However, because of the virus’s long incubation period, the high proportion of infected individuals that are asymptomatic,^6^ and the long period in which apparently-recovered individuals can continue to shed virus in their stool,^7–10^ screening FMT donors using COVID-19 clinical assessment alone is insufficient.

Despite the consensus that FMT donors should be screened for SARS-CoV-2, the optimal available strategy for detecting asymptomatic carriage among FMT donors is unclear. The theoretical effectiveness of polymerase chain reaction (PCR) tests using nasopharyngeal swabs, stool-based PCR tests, donor serology tests, or a combination of those tests has not been assessed or compared. We therefore developed a mathematical model of SARS-CoV-2 infection among FMT donors that simulates the effect of different testing strategies. The model quantifies the effect of more stringent testing on the desirable reduction in potentially infectious, virus-positive donations processed into FMT material and released for use as well as the undesirable reduction in virus-negative donations released.

## METHODS

We built an abstract model of FMT donors, simulating their donation schedule, SARS-CoV-2 infection incidence, and COVID-19 disease course. On top of these simulations, we layered various screening strategies, accounting for the imperfect specificity and sensitivity of each test, to estimate how many virus-negative donations would be appropriately released for use and how many virus-positive donations would be undesirably released. Parameters for the model are shown in Table 1.

**Table 1.** Parameter values. See Supplemental Table 1 for a summary of the meanings of the *I*_1_, *I*_2_, *R*_1_, and *R*_2_ categories.

| Parameter | Point estimate | Lower bound | Upper bound | Source |
| --- | --- | --- | --- | --- |
| Maximum simulation length (days) | 365 | — | — | Assumption<sup>a</sup> |
| Days between donations | 4 | 1 | 10 | Assumption<sup>a</sup> |
| Incidence (daily probability of infection) | $10^{-4}$ | $10^{-5}$ | $10^{-3}$ | Assumption<sup>b</sup> |
| $I_1$ duration (days) | 2 | 1 | 4 | 8,9,15,16 |
| $I_2$ duration (days) | 5 | 3 | 10 | 8,9,15,16 |
| $R_1$ duration (days) | 7 | 3 | 15 | 8,9,15–18 |
| Probability that an infected donor is symptomatic | 0.35 | 0.20 | 0.50 | <sup>6</sup> |
| Probability that an infected donor sheds virus in stool | 0.5 | 0.33 | 0.66 | 7,10,15,16 |
| Days of donations rejected prior to development of symptoms | 14 | — | — | Assumption<sup>a</sup> |
| Days between serology tests | 60 | — | — | Assumption<sup>a</sup> |
| Serology test sensitivity | 0.95 | 0.8 | 0.99 | <sup>19</sup> |
| Serology test specificity | 0.95 | 0.8 | 0.99 | Assumption |
| Days between swab tests | 14 | — | — | Assumption<sup>a</sup> |
| Swab test sensitivity | 0.75 | 0.5 | 0.95 | <sup>20</sup> |
| Swab test specificity | 0.99 | 0.95 | 1 | Assumption |
| Stool test sensitivity | 0.9 | 0.5 | 0.99 | Assumption<sup>c</sup> |
| Stool test specificity | 0.99 | 0.95 | 1 | Assumption<sup>c</sup> |

a: Informed by operations at a large stool bank.
b: The point estimate of $10^{-4}$ corresponds to 35,000 daily cases in a population of 350 million, approximating the US average in early April 2020.
c: Informed by an assay being implemented at a large stool bank.

Each simulation treats a single donor and runs in discrete time steps of 1 day. The simulation begins when the donor enrolls as a stool donor and ends either when the donor is removed because of a positive virus test or after a fixed time. Donors donate stool at set intervals.

We simulate the course of SARS-CoV-2 infection according to the general picture of Sethuraman *et al*.^13^ For simplicity, and to reflect the rigor of the initial screening for new donors, donors are assumed to have tested negative on all screens be unexposed *U* when they enroll on the first day of the simulation. Each day, the donor has a probability of becoming infected *I*_1_; this is the incidence of infection. We ignore any latent period, as it is not relevant to the model. A proportion of infected donors develop symptoms. If a donor becomes symptomatic, their donations from the 14 days prior to onset of symptoms are rejected, and the donor is removed. We assume that an asymptomatic donor in phase *I*_1_ has detectable virus in their nasopharynx but is not shedding detectable loads virus in stool and has not developed detectable IgG antibodies (Supplemental Table 1). After a period of time, the donor enters a second phase of infection *I*_2_. A proportion of *I*_2_ donors are “stool shedders”. Shedders have detectable virus in their stool, and donations produced by shedders are virus-positive. After this second phase of infection, donors enter a first recovery phase *R*_1_. In this phase, donors no longer have detectable nasopharyngeal virus, but they do have detectable antibodies. Shedders continue to produce virus in stool. Finally, donors enter a second recovery phase *R*_2_. Donors in this phase do not have detectable virus in their stool, but they remain detectable by serology. We did not consider the role of immunity because the chance of multiple asymptomatic, undetected infections during the simulation period is low.

Simulated donors are screened for the virus according to a screening strategy that consists of a set of individual test types: a nasopharyngeal swab test performed at 14-day intervals; a blood IgG antibody test at 60-day intervals; and a stool test performed at 14-day intervals, at 28-day intervals, or at every donation. If a donor tests positive on any test implemented in a strategy, they are removed and do not continue to donate. Donations are released only if they are “bookended” by two negative screens. In other words, any donations made after the last negative test conducted before the first positive test are destroyed.

The model has two outcomes: the number of “true negative”, virus-negative donations released and the number of “false negative”, virus-positive donations released. A desirable screening strategy will release many virus-negative donations and few or no virus-positive donations, while a poor strategy will needlessly destroy many virus-negative donations or release many virus-positive donations.

To evaluate the effectiveness of different testing strategies, 10,000 simulations were run for each of 3 incidences (1 infection per 1,000 people per day; 1 per 10,000; 1 per 100,000) and each of 9 screening strategies (stool testing only at 28-day intervals or 14-day intervals, or testing every stool; nasopharyngeal swabs only; nasopharyngeal swabs and stool at each of the 3 stool-testing intervals; nasopharyngeal swabs and serology; nasopharyngeal swabs, serology, and every stool). A sensitivity analysis was run to evaluate the dependence of the model outcomes on input parameters. In 10,000 simulations, parameters were varied over the hypercube bounded by the upper and lower parameter estimates in Table 1. Sensitivity was assessed by the Spearman’s *ρ* correlation between each input parameter and each of the 2 outcomes. Statistical significance was assessed using the false discovery rate, treating the simulations in each strategy separately.

Simulations and analyses were run using R (version 3.6.0).^14^ Code to reproduce the results is available online (DOI: 10.5281/zenodo.3903840).

## RESULTS

The number of virus-positive and -negative donations released varied over simulations and depended on testing strategy and incidence of infection (Figure 1, Supplemental Table 2). In general, the more sensitive strategies released fewer virus-positive donations but also removed donors early due to false positives and therefore released fewer virus-negative donations per donor. In other words, the most sensitive strategies were also the least specific.

**Figure 1.**
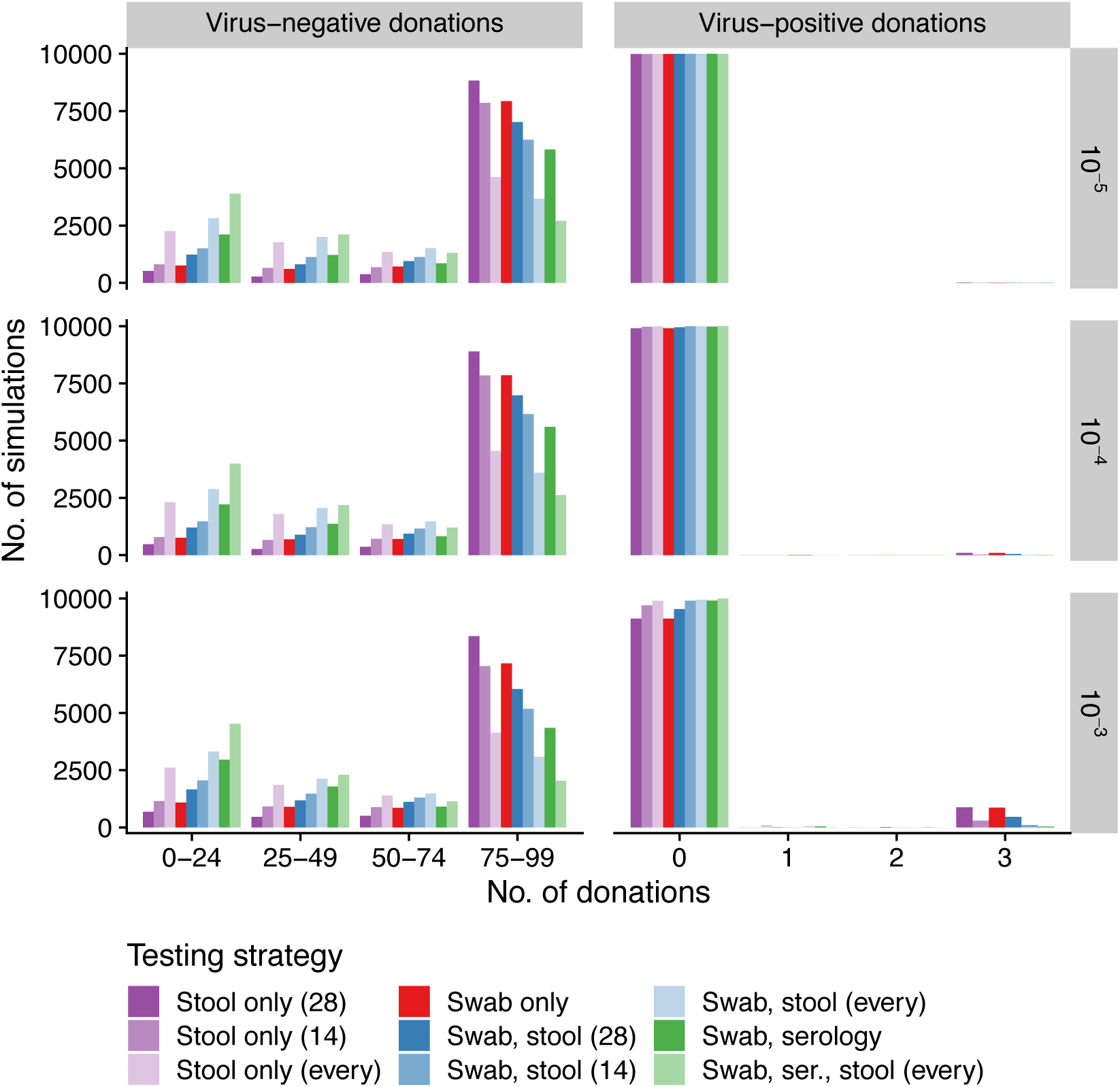
Number of virus-negative and -positive donations released (columns, *x*-axis) across simulations (*y*-axis) for different daily incidences (rows, infections per person per day) when using different testing strategies (colors). Swabs are always at 14-day intervals and serology is always at 60-day intervals. Stool tests are performed at 14-day intervals, 28-day intervals, or for every donation.

At the baseline incidence of 1 infection per 10,000 people per day, the least stringent strategy (testing stool at 28-day intervals) released approximately 1 virus positive-donation per 3,000 donations, while the most stringent strategy (nasopharyngeal swabs, serology tests, and testing every stool) released approximately 1 per 400,000 (Supplemental Table 2). In other words, the most stringent strategy released 100-fold less virus-positive material than the least stringent strategy.

Even at the lower incidence of 1 infection per 100,000 people per day, the least stringent strategy (testing stool at 28-day intervals) released approximately 1 virus-positive donation per 30,000 donations. By contrast, the most stringent strategy (nasopharyngeal swabs, serology tests, and testing every stool) released 1 virus-positive donation per 40,000 donations only at the higher incidence of 1 infection per 1,000 people per day. In other words, the difference in risk of released virus-positive material between the most and least stringent strategies was comparable to the effect of a 100-fold change in daily SARS-CoV-2 incidence.

In a sensitivity analysis (Supplemental Figure 1), the parameters most strongly associated with the two outcomes (Spearman’s ρ > 10%, false discovery rate < 0.05) were donation interval (longer interval correlated with fewer virus-negative donations released), the specificities of the 3 tests (more specific tests correlated with more virus-negative donations released), and SARS-CoV-2 incidence (higher incidence correlated with more virus-positive donations released).

## DISCUSSION

A mathematical model of SARS-CoV-2 infection among stool donors suggests that, if incidence among stool donors is comparable to the aggregate national average, if a stringent strategy is used, and if our estimates of the sensitivity and specificity of the tests are accurate, then the probability of releasing a virus-positive donation for clinical use is low. The most stringent test strategies involved testing every stool, while the least sensitive strategies were to use nasopharyngeal swab alone or to test stool at 28-day intervals. More stringent tests were more sensitive but also less specific, and the most appropriate strategy must be determined by a balance between the necessary stringency and logistical considerations like resourcing.

The strength of this analysis is its quantitative treatment of a pressing clinical question. However, it has multiple limitations. First, as a modeling study, the accuracy of the results depend on the accuracy of the input parameters and the appropriateness of the model structure, especially the tests’ sensitivity and specificity as well as the incidence of SARS-CoV-2 infection, values which remain subject to refinement. Thus, the quantitative predictions made by the model should be used as guides to clinical reasoning rather than as precision forecasts. Second, the model makes a number of assumptions about the course of disease that may be shown to be invalid or that are no longer applicable. For example, our assumption that newly enrolled donors are seronegative maximizes the sensitivity of serology testing. As the number of candidate donors with positive serology rises, the sensitivity and utility of the serology test will decline. Finally, verifying the model would be challenging, as the possibility of fecal-oral transmission of SARS-CoV-2 has not been confirmed, and there is no accepted “gold standard” for detecting SARS-CoV-2 in stool.

Although these results are encouraging, we again caution that they depend on a number of assumptions about testing quality and SARS-CoV-2 epidemiology that will be refined in the coming months. Nevertheless, this method is valuable in assessing the risks of transmission in this evolving pandemic, and we hope this approach can serve as a model for evaluating testing strategies for other pathogens or human-derived therapies beyond FMT.

## Acknowledgements

Emily Langner for helpful comments.

---

**Supplemental Table 1.** Summary of donor disease course phases and their relationship to each test.

| Status | Signs/<br>symptoms | Nasopharyngeal<br>test | Stool<br>test | Serology<br>test |
| --- | --- | --- | --- | --- |
| Unexposed ($U$) | – | – | – | – |
| Infected but not<br>shedding ($I_1$) | + (if<br>symptomatic) | + | – | – |
| Infected and<br>shedding ($I_2$) | + (if<br>symptomatic) | + | + (if a<br>shedder) | – |
| Recovered but<br>shedding ($R_1$) | – | – | + (if a<br>shedder) | + |
| Fully recovered<br>($R_2$) | – | – | – | + |

**Supplemental Table 2.** Virus-positive donations released over 10,000 simulations. Values shown are the proportion of total released donations that are virus-positive, 95% confidence interval on that proportion, and the raw counts of virus-positive donations / total released donations. Proportions are shown to one significant digit.

| Testing strategy | Daily incidence (infections per person per day) |  |  |
| --- | --- | --- | --- |
| | $10^{-5}$ | $10^{-4}$ | $10^{-3}$ |
| Stool only (28) | 4 pcm (3 pcm to 5 pcm; 33/848185) | 30 pcm (30 pcm to 40 pcm; 288/851793) | 300 pcm (300 pcm to 300 pcm; 2636/819180) |
| Stool only (14) | 2 pcm (1 pcm to 4 pcm; 18/793167) | 10 pcm (8 pcm to 10 pcm; 77/793966) | 100 pcm (100 pcm to 100 pcm; 901/743270) |
| Stool only (every) | 0 pcm (0 pcm to 0.6 pcm; 0/593511) | 2 pcm (1 pcm to 4 pcm; 12/585989) | 20 pcm (20 pcm to 20 pcm; 106/556362) |
| Swab only | 3 pcm (2 pcm to 4 pcm; 21/799372) | 40 pcm (30 pcm to 40 pcm; 284/794911) | 300 pcm (300 pcm to 400 pcm; 2617/751393) |
| Swab, stool (28) | 1 pcm (0.6 pcm to 2 pcm; 9/739826) | 20 pcm (20 pcm to 20 pcm; 153/737824) | 200 pcm (200 pcm to 200 pcm; 1394/677216) |
| Swab, stool (14) | 0.9 pcm (0.3 pcm to 2 pcm; 6/693695) | 4 pcm (2 pcm to 6 pcm; 26/690734) | 50 pcm (40 pcm to 50 pcm; 306/623760) |
| Swab, stool (every) | 0 pcm (0 pcm to 0.7 pcm; 0/528625) | 1 pcm (0.5 pcm to 3 pcm; 7/519153) | 10 pcm (10 pcm to 20 pcm; 61/479578) |
| Swab, serology | 0.5 pcm (0.1 pcm to 1 pcm; 3/635133) | 6 pcm (4 pcm to 9 pcm; 39/618887) | 30 pcm (30 pcm to 40 pcm; 177/530406) |
| Swab, ser., stool (every) | 0 pcm (0 pcm to 0.9 pcm; 0/430492) | 0.2 pcm (0.006 pcm to 1 pcm; 1/418555) | 3 pcm (1 pcm to 5 pcm; 10/369524) |

pcm = per cent mille = per 100,000

**Supplemental Figure.**
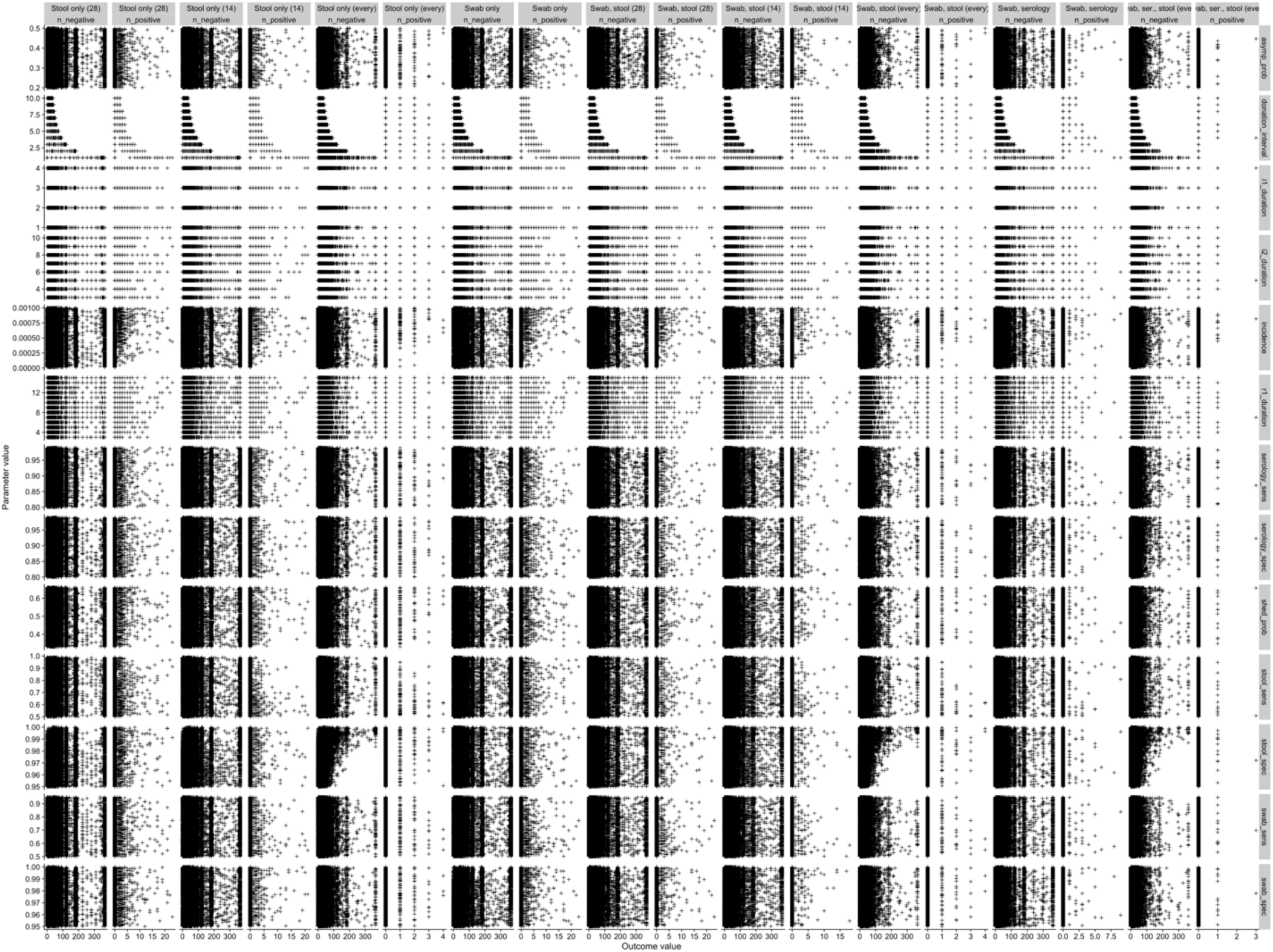
Sensitivity analysis. Each point represents 1 simulation. Columns show testing strategies and outcomes (number of donations released). Rows show input parameters.

